# Decomposing Predictive Information in Social Dynamics

**DOI:** 10.1101/2025.05.16.654393

**Authors:** Akira Kawano, Liam O’Shaughnessy, Radmila Neiman, Kosmas Deligkaris, Luis Carretero Rodriguez, Ichiro Masai, Greg J. Stephens

## Abstract

Social behaviors include some of the most interesting interactions in living systems yet their principled characterization remains unsolved. Here we suggest that at the core of social interactions is the notion of mutual prediction, which we analyze in the context of two male zebrafish engaged in a dominance contest. Using 3D velocity trajectories, we construct the mutual information between a two-animal past and one-animal future, and we quantify the redundant, unique, and synergistic components using partial information decomposition across time windows. We find that predictive information decomposition naturally aligns with important social concepts, such as mirroring and dominance. At contest end, we find asymmetries in self-unique and redundant information that reflect the emergent dominance relationship. Applied to *mecp2* zebrafish mutants, an autism model, we find that predictive information is reduced overall, but especially for synergistic flows, which is indicative of difficulties in more complex social dynamics.

**Significance Statement:** Social interactions are rich and diverse, ranging from mirroring to complementary actions. A unifying framework for defining and analyzing such interaction types has long been needed. Here, based on modern information theory, we formulate how the past state of interacting organisms encodes the future state of an individual. This framework provides a natural decomposition of pairwise social dynamics into mirroring, independent action, directed influence, and joint action. Applied to dominance contests in zebrafish, these modes of interaction capture distinct phases of conflict, their assessment strategies, and the resulting dominance relationships. Moreover, our analysis reveals a specific disruption in the social behavior of mutant zebrafish linked to autism, shedding new light on impairments in communication and social learning.

## Introduction

Resulting from interactions between individuals, social behavior occurs both among and across species, from microorganisms to humans. These interactions are broadly important; cooperation is thought to have played a crucial role in the evolution of multicellularity [1], and human sociality is a primary determinant of mental and physical well-being (see e.g. [2]). Despite such universal importance and in part due to it’s complexity, social behavior has been predominantly analyzed either qualitatively or with specifically crafted measures and experiments. These include the frequency and context of human-derived action patterns, as in an ethogram, and discipline-defined experimental protocols, as in social neuroscience (see e.g. [3]). Another approach is to account directly for the interactions through modeling, for example, by viewing the social system as a collection of interacting particles [4]. While this later approach is promisingly general, interactions can change abruptly in time, making estimation difficult. Also, the form of the interactions varies with the choice of behavioral coordinates, which can hinder interpretation.

An alternative view is to focus on social dynamics as a reciprocal flow of influence, with intertwined actions and reactions [5, 6]. Such an approach naturally aligns with the broad perspective of social behavior as communication; we interact to change and be changed by our interaction partners [7]. But how do we quantify the volleys of dynamics within this reciprocal perspective? Here we introduce an information-theoretic framework, which combines partial information decomposition (PID) [8] with prediction, to characterize the flows of information between participants in a social system. Our approach leverages the representation invariance of information theory and is thus less dependent both on human-derived measures and on particular movement trajectories, which can vary in presentation across organisms even in similar social contexts. As PID provides a fundamental decomposition of information flow, we also generalize previous approaches that used transfer entropy to determine dominance [9]. We compute and interpret these information flows using both previous work on the pairwise dominance behavior of adult zebrafish [10], as well as new behavior measurements of *mecp2* zebrafish mutants, which have been linked to Autism Spectrum Disorder [11].

### Decomposing predictive information in social dynamics

As a fundamental framework for understanding social dynamics, we focus on the concept of mutual prediction in the simplest social case of two individuals, Fig. 1(A). Our perspective is motivated by growing empirical evidence of social coupling, wherein the brain encodes information about an interacting partner [12, 13]. Accordingly, we aim to describe how the joint past state of the two individuals 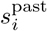 and 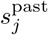 predicts the future state of either 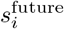. This predictive relationship is naturally quantified using the joint mutual information,

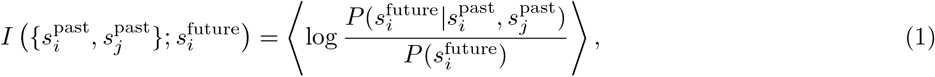

**FIG. 1:**
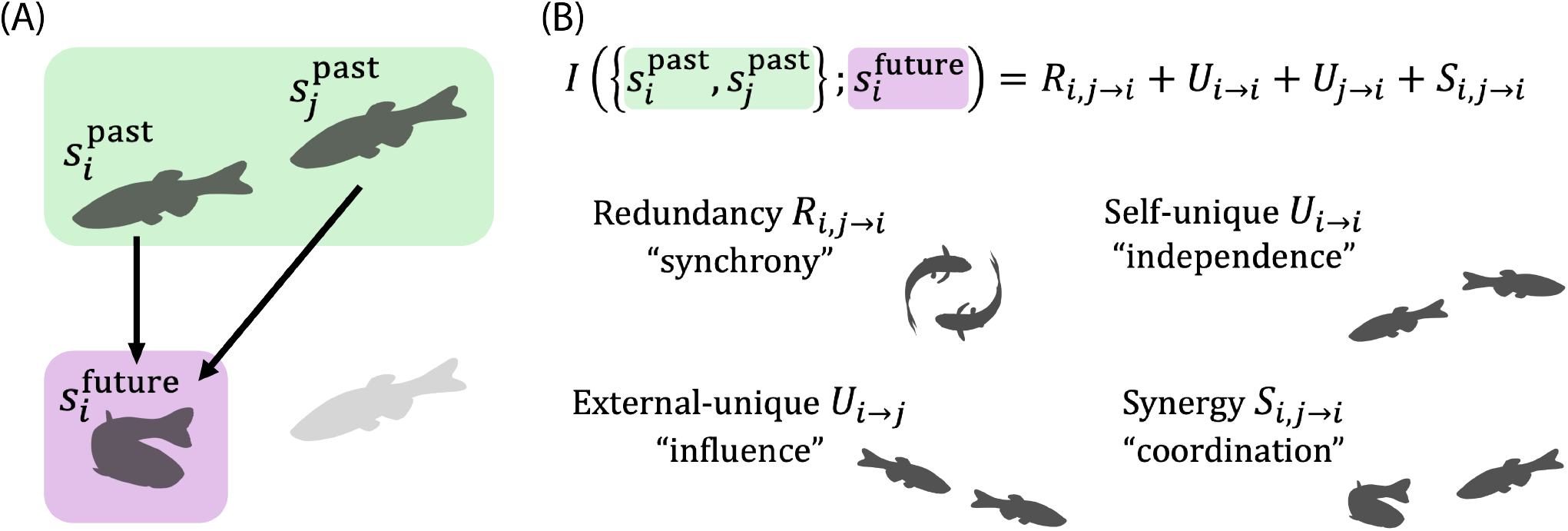
Decomposing predictive information in social dynamics. (A) In social behavior, the future state of a focal individual 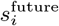 depends on both the past state of itself 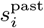 and the past state of others 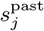. Even the minimal social system of two agents thus involves three variables that support multiple modes of interactions, such as mirroring and complementary actions between individuals. (B) We introduce partial information decomposition (PID) of predictive information for social dynamics. In PID we decompose the total predictive information 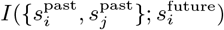 into four information terms, where redundant information *R*_*i,j*→*i*_, self-unique information *U*_*i*→*i*_, external-unique information *U*_*j*→*i*_, and synergistic information *S*_*i,j*→*i*_ quantifies the degree of synchrony, independence, directional influence, and joint coordination, respectively.

where ⟨*· · ·* ⟩ denotes an average over the joint distribution, 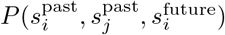. We note that this predictive information is a function of three variables and thus generally includes higher-order interactions [14–16]. The mutual information of Eq. 1 is a principled, scalar measure of the average amount of instantaneous predictive information. Yet, this overall information can result from a variety of processes. For example, how much of the future depends on the state of both individuals together? How much of the future of one individual is determined by itself or by the other? Previous research has examined directed influence using both Granger causality and it’s nonlinear generalization, transfer entropy [17]. However, the framework of partial information decomposition (PID) [8, 18, 19], from which transfer entropy can also be computed, provides a more elemental picture. We argue here that PID of predictive information is a natural and quantitative architecture for capturing information flow resulting from social interactions, and thus for achieving deeper understanding of social behavior.

In PID we decompose the total or multi-information of a collection of variables into a sum of non-negative “information atoms” that capture redundant, synergistic and unique information present within the system. In an example with three variables {*A, B, C*} the redundant information that {*A, B*} carry about *C* is information this is contained in either *A or B*, while synergistic information is information that {*A, B*} carry about *C* that requires both *A and B*. The unique information that *A* carries about *C* is information about *C* that is not contained in *B* and similarly for the unique information that *B* carries about *C*. Synergy and redundancy have been used to characterize neural activity [20] and large-scale human brain dynamics [21], and PID has also been used to understand gene expression [22].

We apply PID to two-organism social dynamics, with the future state of the focal organism 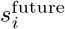 as the target and the past states of both organisms, 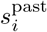 and 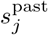 as predictors Fig. 1(B),

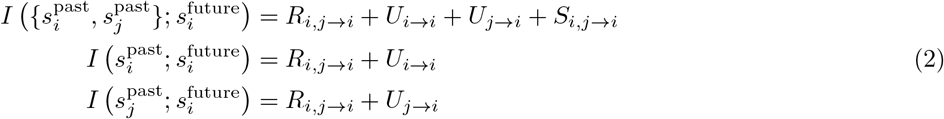

where *R*_*i,j*→*i*_ denotes the redundant information about 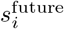 shared by 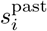 and 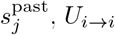 and *U*_*j*→*i*_ are unique information from 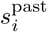 and 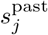 respectively about 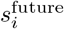, and *S*_*i,j*→*i*_ represents the synergistic information about 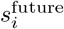 available only in the joint state,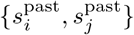. The redundant information *R*_*i,j*→*i*_ is the degree to which predictive information is shared [23] and captures mirroring or synchronization between organisms. The self-unique information *U*_*i*→*i*_ is the information isolated within the focal organism, thus providing a measure of independence: organisms with larger self-unique information are more autonomous. In contrast, the other-unique information *U*_*j*→*i*_ is the information transferred from the other to the focal organism and thus characterizes directional influence. The synergistic information *S*_*i,j*→*i*_ is the predictive information modified by the opponent, capturing joint actions or coordination. An example of such coordination is the division of labor, as seen in leader–follower roles. Leaders determine the future group state, while followers adapt to the leader’s actions, where the followers’ predictive information is modified by the behavior of the leaders. While we introduce information flow in the context of two organisms, our approach can also be applied to larger groups, a point we discuss further in the discussion. Finally, we note that the transfer entropy is determined explicitly in terms of information atoms [24], 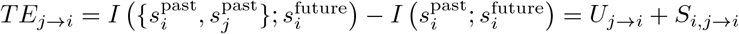.

### Joint coordinates and interaction changes across behavioral epochs

We explore our framework of predictive information flow in the context of the social behavior of zebrafish, an important vertebrate genetic model system. We focus on a pair social setting where two previously isolated adult male zebrafish are introduced into a novel environment, in which they then seek to establish a dominance hierarchy [25, 26]. We track their movements at high spatiotemporal resolution using a single 3D coordinate for each fish, the “pectoral point” located approximately at the base of the pectoral fins. Both the experiments and tracking have been previously described [10].

We quantify the movement trajectory of each fish using a suitably determined 3D velocity, Fig. 2(A). For each individual, we introduce a co-moving coordinate system which is spanned by the axis from the focal to the other organism 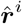, the lab-frame *z* axis 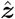, and a 3rd orthogonal axis 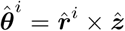. We compute the pectoral point velocity ***V****^i^* using finite differences and then project onto the co-moving frame so that 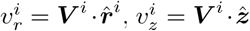, and 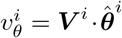. We note that our coordinate system allows for ready interpretation. For example, the approach velocity *v_r_* captures important agonistic interactions such as attack and chasing behavior, where *v_r_* is predominantly positive for the chaser and negative for the fish being chased, Fig. 2(B). We focus our following analysis on *v_r_*, with the corresponding results for *v_θ_* and *v_z_* provided in SI.

**FIG. 2:**
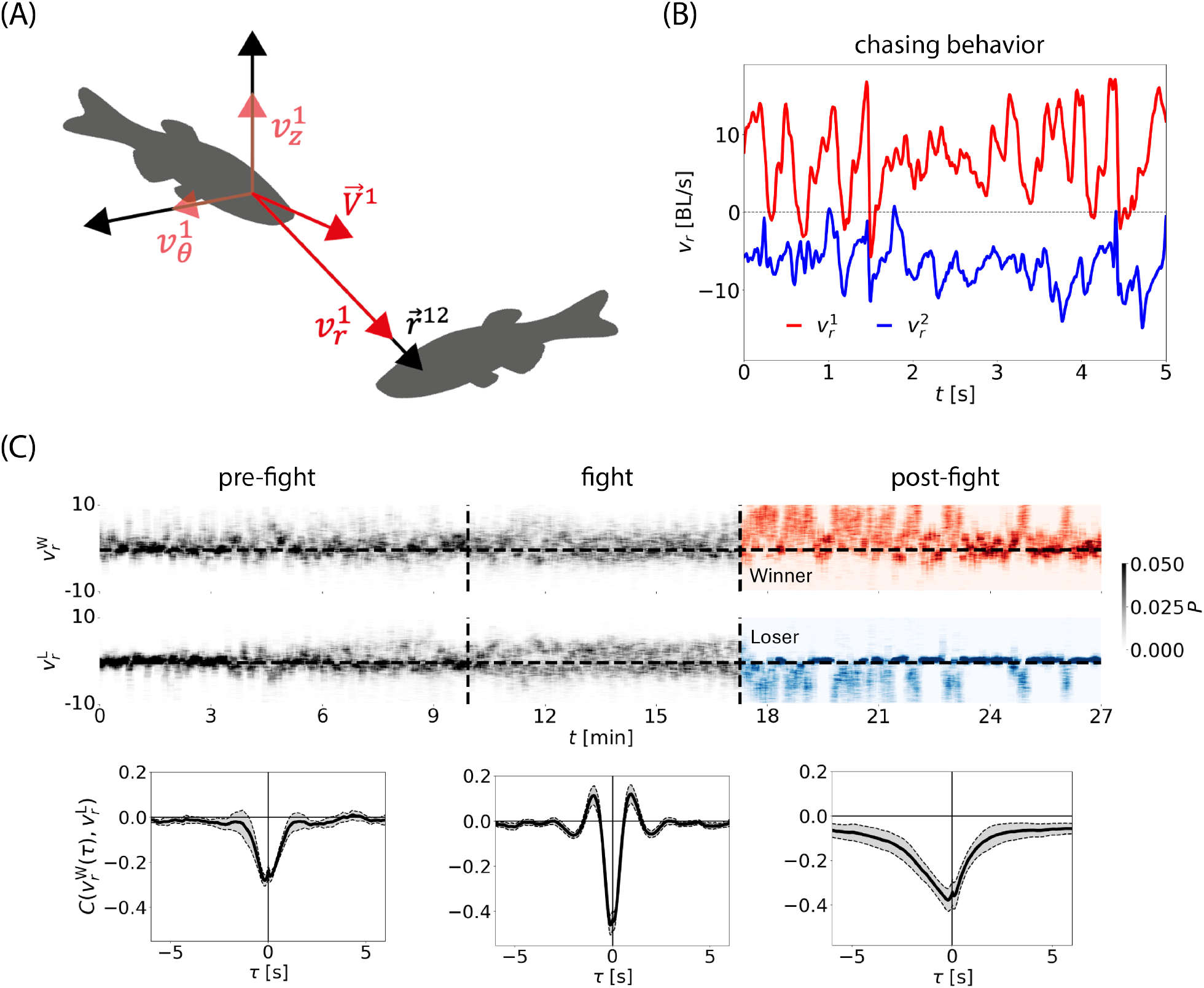
We introduce co-moving coordinates and show that interactions change dramatically across behavioral epochs. (A) We capture interactions through movements using projections of the velocity vector ***V***^1^ onto the axis from the focal individual to another 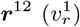, onto the *z*-axis 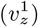, and the axis orthogonal to the other two axes 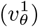. (B) *v_r_*reflects approaching movements, such as chasing behavior, and we show an example time series where fish 1 chases fish 2. (C) Social dynamics of two male zebrafish show sequence structure, i.e., epochs, consisting of “pre-fight”, “fight”, and “post-fight”. The approach velocity *v_r_*is peaked around *v_r_*= 0 in pre-fight epochs while it is more varied during the fight. In post-fight epochs, the approach velocity of the winner 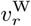 tends to be positive while the approach velocity of the loser 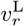 is generally negative, reflecting the predominance of winners chasing losers. We compute the average cross-correlation function of *v_r_*between the (eventual) winner and loser fish 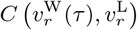 (Methods: Cross-correlation function) in each epoch across experiments. In pre-fight epochs, the magnitude of the correlation, i.e., coupling, is small. In fight epochs the coupling is larger and displays oscillations reflecting cyclic attacks. In post-fight epochs, the time-symmetry is broken, demonstrating larger influences from winner to loser. We use predictive information decomposition to systematically describe such changes in the interactions.

An important feature of social interactions is that they can change abruptly as information is transferred between participants; just imagine the reaction when walking into a sports bar and loudly rooting for the non-local team! In our two-fish setting of a dominance contest, these changes can be coarsely characterized as behavioral epochs that are quantitatively distinguished by the nature of the movements [10]: a pre-fight epoch with relatively weak interactions and symmetry between individuals, a fight epoch with strong interactions, and a post-fight epoch with a clear between individual symmetry breaking, Fig. 2(C, top). We use behavior in the post-fight epoch to designate the individual fish as “winner” or “loser”. Among other attributes, the winner does most of the chasing.

We illustrate the interactions within each epoch though the cross-correlation function 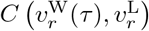, Fig. 2(C) (Methods: Cross-correlation function). In the pre-fight epoch, the overall magnitude of the correlation, the linear coupling between individuals, is small. In the fight epoch, the larger coupling with an oscillation reflects cyclic attacks, where a fish being attacked will attack back after approximately 1 s. The symmetry of the interactions disappears after the fight with larger correlations for a negative time shift, indicating a larger influence from winners to losers. To systematically understand such changes of interaction, we next apply our predictive information decomposition to the dynamics of *v_r_*.

### Predictive information flows within behavioral epochs

We define a one-step future and a *τ* -step past for each organism, Fig. 3(A),

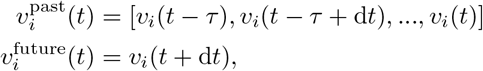

and we illustrate our approach using the approach velocity *v^i^*. We choose *τ* = 5 s to account for the significant crosscorrelations, and we estimate PID using sliding time windows across the recording and a Gaussian approximation for the full distribution. Sliding time windows allow us to account for changing interactions Fig. 3(B-left) and we choose the window size to maximize the total information flow, Fig. 3(B-right). Note that this window size Δ*t*^∗^ = 30 s is substantially longer than the past length *τ*. In the Gaussian approximation both the full predictive information of Eq. 1 and the partial information decomposition of Eq. 2 are analytically computable (Methods: Partial information decomposition for Gaussian variables). We provide movies of example social behaviors showing characteristic information flows in SI.

**FIG. 3:**
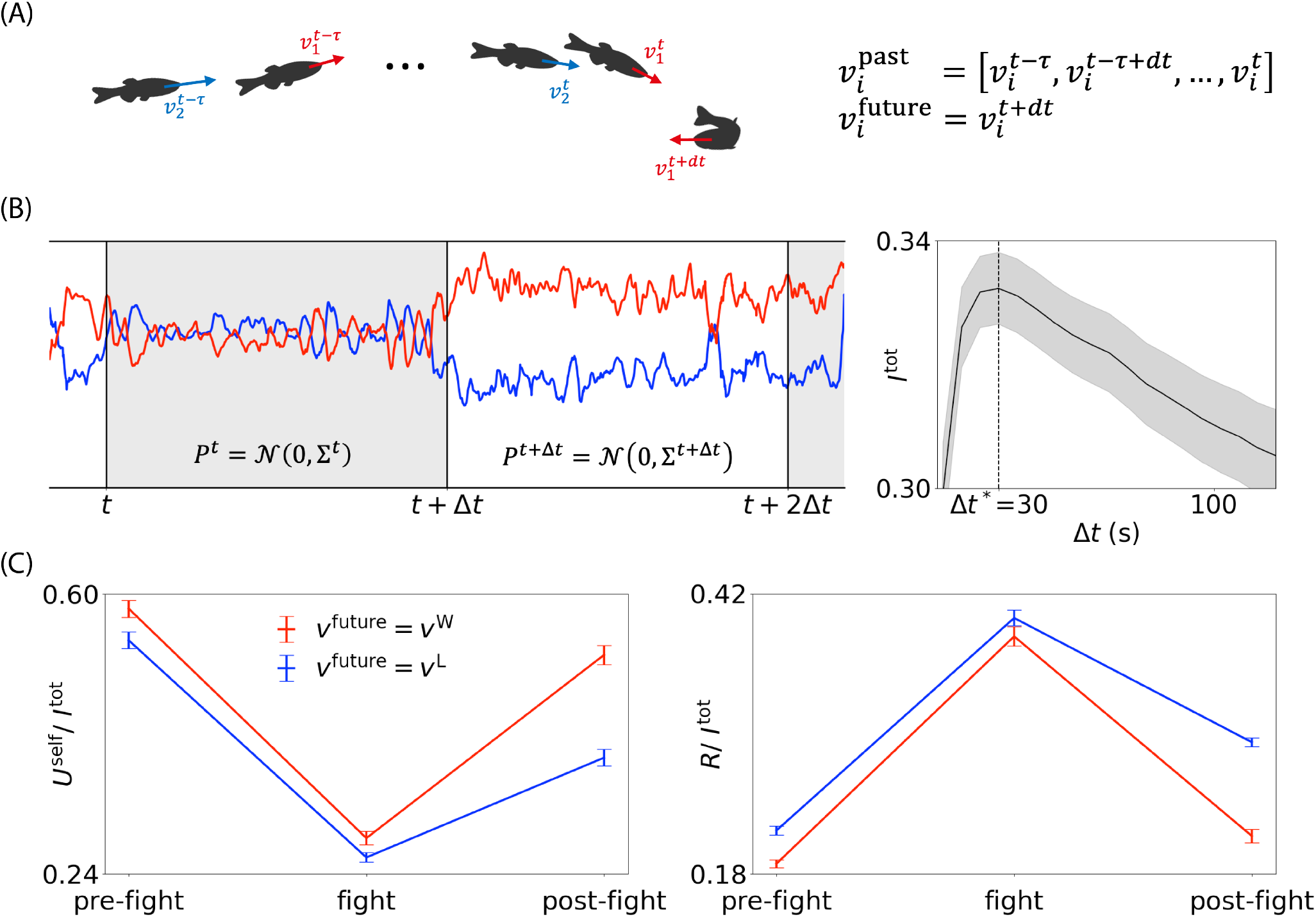
Predictive information decomposition within epochs. (A) We define a one-step future and a *τ* -step past for each organism, and set *τ* = 5 s to include significant temporal correlations, see Fig. 2(C). (B) We estimate predictive information decomposition using sliding time windows across the recording, which captures changing interactions, and a Gaussian approximation for the full distribution, for which an analytic PID expression is available. We choose the window size Δ*t*^∗^ = 30 s to maximize the total information flow 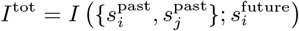 in the Gaussian limit, Eq. 3. (C) We normalize the partial informations by *I*^tot^ in each window and compute the mean information within each epoch. Plots show the average across experiments of the mean information, and error bars show SEM. Self-unique information is high and symmetric in the pre-fight epoch, reflecting more independent movements. Redundancy increases during the fight, suggesting more coupled dynamics. Self-unique information rises again in the post-fight epoch, with an asymmetry reflecting the emergent dominance relationship, demonstrating that the winning organism is more independent.

For each of the behavioral epochs (pre-fight, fight, post-fight) from each experiment we compute the predictive partial information decomposition of the approach velocities averaged across windows and normalized by the total information flow. We then average across experiments and show the results in Fig. 3(C). We find that self-unique information is high with the pre-fight epoch, implying that the fish are more independent, while redundant information increases during fighting, reflecting more synchronized movements. Self-unique information rises again in the post-fight epoch and includes a notable between-organism asymmetry: the self-unique information of the dominant fish is higher. Thus one aspect of the emergent hierarchy is that the movements of the dominant fish are more independent. Redundant predictive information is higher for losers, reflecting their tendency for more mirroring. Interestingly, this is in agreement with results from neural recordings of fighting mice, where subordinate’s neurons predict opponent behavior more than dominant’s neurons [13]. Synergy follows the same trend of redundancy, high information during the fighting epoch and a post-fight asymmetry with higher information in losers than in winners, Fig. S2. This indicates that, fish coordinate their movement with an opponent more in fight, and in post-fight losers respond more to opponent movements compared to winners. Finally, we note that opponent-unique information is approximately zero across all epochs, Fig. S2, indicating that components of future movement uniquely determined by the opponent are negligible. This is essentially due to the larger auto-correlation compared to cross-correlation (Methods: Relationship between unique information and correlation functions). We note that a multidimensional extension of the target, beyond a one-step future, could reveal non-zero opponent-unique information, as further explored in Discussion.

### Predictive information flows across behavioral epochs

To further understand how predictive information flows change during the span of our recordings, we align the window average to the start and stop of each contest, which was determined previously [10], and find that redundant, unique, and synergistic information flows display distinct dynamics across epoch boundaries, Fig. 4. Redundancy ramps up from the start of fight until the end, while self-unique information continuously decreases until the end of fight, suggesting that fish are increasingly engaged in interaction until the winner-loser decision. Interestingly, synergy shows a different trend than redundancy, remaining roughly constant for the duration of the fight. This constant synergy implies that a specific type of information exchange through their joint actions occurs at a constant frequency. In animal contest studies, the information exchange at constant frequency is considered a constraint from the mutual assessment process in which animals assess their opponent’s ability relative to their own [27].

**FIG. 4:**
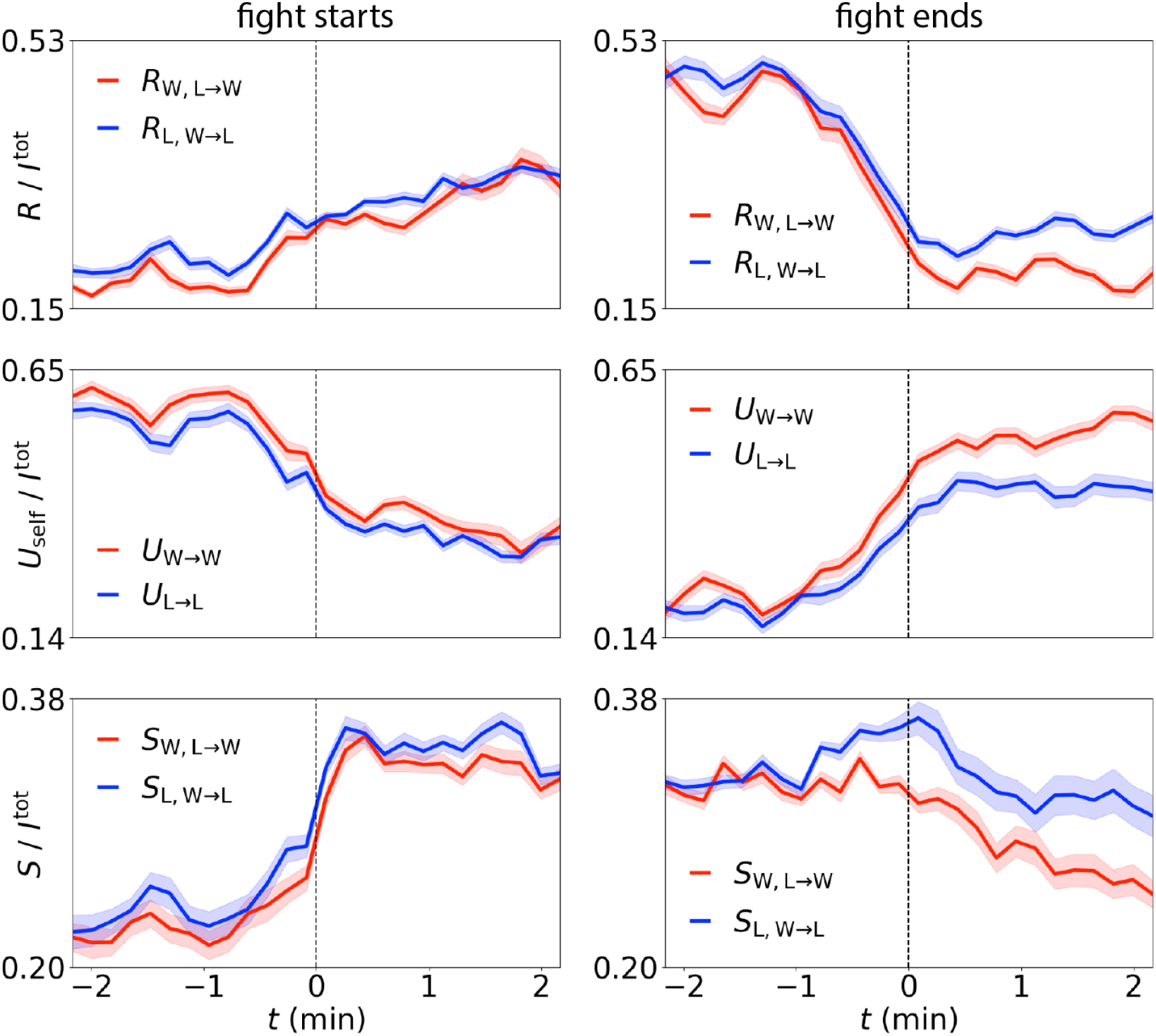
Redundant, unique, and synergistic information flows display different dynamics across contest boundaries. (Left) We align by the start of the fight epoch and compute the average and SEM of the fraction of partial information across experiments. (Right) We align by the end of the fight epoch. Redundancy increases from the start of the fight until the end, while self-unique information continuously decreases until the end of the fight. Synergy shows a different trend than redundancy, remaining roughly constant for the duration of the fight. There are clear post-fight asymmetries across all flows.

### Predictive information flows are disrupted in social mutants

The *mecp2* mutant zebrafish is a model of Autism Spectrum Disorder [28] and displays clear behavioral abnormalities including motor anomalies and defective thigmotaxis [29]. We capture disruptions in mutants’ social interaction by comparing predictive information flows between wildtype zebrafish pairs and *mecp2* mutant pairs from 30 s windows across a 5-hour recording. The total information flow 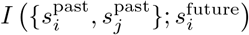) is lower in mutants than in wildtype, as shown in Fig. 5(A). This result suggests that the social behavior of mutants is less structured and less predictable compared to that of wild-type, potentially reflecting reduced motor activity [29], impaired sociability, or a combination of both factors. We then compute the average of the normalized partial informations, Fig. 5(B). Mutants show higher unique information and lower synergy, suggesting more independence and less ability to coordinate in social situations, which aligns with expectations for an autism model [28]. Interestingly, redundant flow is similar to wild type, suggesting that behavioral mirroring, which may require less cognitive demand than complementary actions, remains intact in this mutant.

**FIG. 5:**
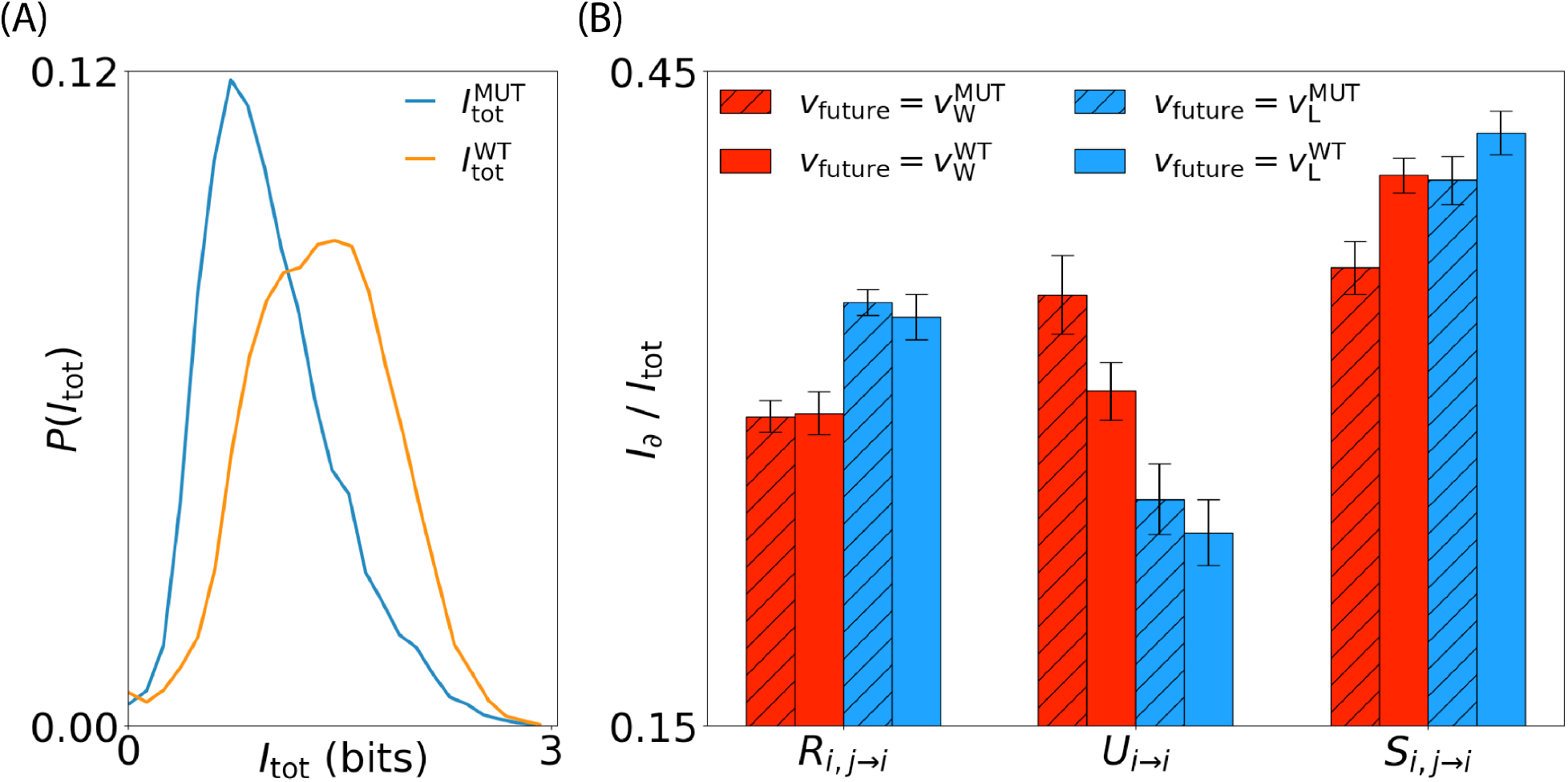
Predictive information flows are disrupted in *mecp2* mutants, a zebrafish model of Autism Spectrum Disorder. (A) The total predictive information flow is lower in the mutants (blue curve) than in wildtype (red curve). We construct these distributions from 30 s windows across a 5-hour recording. (B) In both mutants and wildtype, the self-unique information flow towards the winner is larger than the flow towards the loser, demonstrating that this flow captures the dominance hierarchy in both cases. In contrast, mutant fish show higher unique information and lower synergy, suggesting more independence and less ability to coordinate in social situations. Interestingly, redundant flow is similar to wildtype, suggesting that behavioral mirroring remains intact in this mutant.

## Discussion

Organisms couple to their environment through an action-perception loop [30, 31], and social interactions are distinguished by the addition of feedback between these loops: the actions of one participant influence the perceptions and actions of another (see e.g. [6]). Here we account for this feedback using predictive information, the mutual information between the past and future of our social system. This predictive information can be expressed in redundant, unique and synergistic components using partial information decomposition. We find that these components are directly interpretable in the example of two zebrafish engaged in dominance interactions, in both wild-type and mutant examples. This interpretability is a key advantage of our information-theoretic framework, distinguishing it from neural network based behavioral clustering methods (e.g., [32]), which systematically classify behavior in an unsupervised manner but can yield clusters that are difficult to interpret.

The generality of our information-theoretic approach, which in principle is agnostic to behavioral coordinates, is ideal for comparative studies, where a key challenge lies in making meaningful cross-species comparisons and moving beyond case studies toward general principles underlying behavioral variation [33]. For example, comparing information flows arising from dominance interactions across species could provide quantitative and complementary insights into the interplay among contest strategies, cognition, and ecology, domains that have traditionally been analyzed through game theory [34–36].

We chose a one-step future *s*^future^(*t*) = *x*(*t* + d*t*) to take advantage of a closed-form solution for PID that is only available for a univariate target. More generally, we can define the future state as the trajectory *s*^future^(*t*) = [*x*(*t* + d*t*)*, x*(*t* + 2d*t*)*, …, x*(*t* + *τ*)], enabling the characterization of higher-dimensional temporal dependencies. In Gaussian variables for example, our partial information decomposition is constructed from all possible pairwise correlations and thus a longer future includes new correlations such as 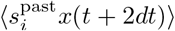. Whether the use of higher-dimensional trajectory future can uncover non-zero opponent-unique information in zebrafish social dynamics is an important direction for future investigation. For a higher-dimensional Gaussian target, a numerical estimator of PID based on the BROJA unique information formulation has been proposed [37]. For generic distributions, a neural network approach based on the BROJA unique information [38] can accommodate high-dimensional targets. As an alternative, we can discretize the time series of *s*^past^(*t*) and *s*^future^(*t*) in any dimension and estimate PID using the original redundancy measure for discrete distributions [8].

Our Gaussian approximation for the joint distribution within each time window enabled the use of a closed-form solution for PID. For general continuous distributions, there are numerical PID estimators based on the Cone Program [39] and neural network optimization [38]. Both are based on BROJA unique information formulation. By discretizing the time-series, we can also estimate PID in discrete distributions based on the redundancy measure using the specific information [8, 22]. By relaxing the Gaussian approximation, we expect to extend the length of time windows and more accurately capture the complexity of social dynamics.

We can naturally extend our approach to more organisms by considering information flow from an N-organism past to a focal-organism future. For example, with a 3-organism past (3 predictors), redundant information about a target will take many forms such as the redundancy between a single predictor *X* and a joint of predictors *Y, Z* and PID offers *m* = 18 modes of information flow. For arbitrary *N*, the decomposition can be determined by the Möbius transform, for which an efficient estimator has been developed [40]. We note however that the number of PID modes grows super-exponentially with *N*, determined by the Dedekind numbers minus two [18], and the interpretation of the resulting very large number PID modes for more than 3-organisms is a significant challenge. We note that it might be possible to coarse grain the information flows into macroscopic flows (see an attempt in ΦID [41]). Another possibility is to focus on the information about the focal individual’s state that is carried by its own past and by the state of all other individuals, 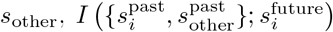, or influence of a focal individual on the dynamics of other,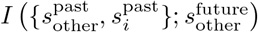. In such formulations, the number of PID modes grows linearly with the number of organisms, making PID analysis more tractable for larger groups, including those exhibiting collective behavior.

For simplicity we focused here on information flow from a two-organism past to a one-organism future. In dyadic settings, we can more generally investigate the information between a two-organism past and two-organism future, using mutual information 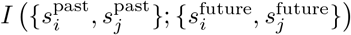. A natural decomposition of such information needs to describe how information encoded in a specific form (unique, redundant, and synergistic) evolves over time and be stored in any of each form in the future. This will introduce a picture of the “product” of two PIDs, one decomposing the information flow from past to future (predictors to targets) and another decomposing the flow from future to past (targets to predictors). Such an idea has been formalized as integrated information decomposition, ΦID [41]. By decomposing information between two predictors and two targets, ΦID provides a larger *N* = 16 set of modes of information corresponding to pairs of original PID modes evolving from past to future. For example, one of the ΦID modes represents information that is uniquely encoded in one of the variables in the past and as redundant information in the future, this can be interpreted as information duplication along the dynamics. Exploring ΦID for social dynamics is a promising future direction.

Zebrafish have emerged as a promising model organism for investigating social dysfunctions, complementary to traditional rodent models [28, 42]. Accordingly, substantial efforts are underway to advance behavioral assays and analysis methods for studying their social deficits using mutants [32, 42]. In this study, we identified behavioral anomalies in *mecp2* mutant zebrafish pairs, characterized by reduced predictive information, particularly in synergistic components, suggesting impairments in more complex social dynamics. We believe that our framework can be a useful tool for detecting subtle behavioral defects in other zebrafish models of neurological disorders, thereby contributing to the systematic investigation of these diseases. In addition, since neuropsychiatric disorders in human are often linked to social dysfunctions [43], our findings highlight the potential of using our framework for identifying biomarkers relevant to psychiatric diagnosis. Such biomarkers related to social dysfunctions may define a new transdiagnostic domain, offering targets for innovative treatments.

## Acknowledgments

This work immediately follows previous research [10] enabled by grant RGP0055 from the Human Frontiers Science Program, and was supported by funds from OIST Graduate University (AK, RN, KD, LCR, IM, GJS) and Vrije Universiteit Amsterdam (LOS,GJS). We thank Tatsuo Izawa from the Biological Physics Theory Unit at OIST Graduate University for useful discussions and zebrafish behavior insight. We are also grateful for the help and support provided by the Engineering and Scientific Computing & Data Analysis sections of Research Support Division at OIST.

## Methods

### Data

We use body point trajectories obtained from a recent advance in the measurement and analysis of zebrafish dominance contests [10], as well as from new experiments for the mutant and wildtype comparison of Fig. 5. A link to the data from Ref. [10] is available in the README file of the following Github project: https://github.com/liamshock/Dynamics of dominance. Briefly, the data consists of the 3D location of three body points (approximately situated near the head, pectoral fins and base of the tail), for each individual of a pair of adult male zebrafish. These body point locations are sampled at 100 Hz across a measurement period which covers pre-contest, contest and post-contest epochs. The dataset includes precise temporal boundaries for fight epochs, and we use *N* = 18 recordings in which a clear dominance structure emerges. For the mutant comparison, social interactions of *N* = 8 wild-type male pairs and *N* = 6 *mecp2* mutant male pairs were recorded for 5 hours at a sampling rate of 140 Hz. The fish were 13–17 months old and the recordings followed 24 to 72 hours of isolation. We analyzed the recordings as described above.

### Zebrafish strains

Zebrafish were maintained at 28.5 *^°^*C on a 14-hour light/10-hour dark cycle, following standard procedures [44]. Okinawa wild-type (oki) was used as a wild-type strain. We used a *mecp2* mutant (sa21196, ZIRC ID ZL11092.02) allele with one point mutation that introduced a premature stop codon, provided by the Zebrafish International Resource Center (ZIRC, Eugene, OR, USA).

### Ethics statement

All zebrafish experiments were carried out under the guidelines of the OIST Animal Care and Use Program, following the principles outlined in the Guide for the Care and Use of Laboratory Animals by the National Research Council of the Nation Academies. These procedures received approval from the Association for Assessment and Accreditation of Laboratory Animal Care (AAALAC). The OIST Institutional Animal Care and Use Committee approved all related experimental protocols (Protocol: ACUP-2023-018, ACUP-2023-022-2).

### Behavioral epochs

The identification of the temporal boundaries of fight behavior was described previously [10]. Based on the boundaries of the fight epoch, we define the pre-fight epoch as the 10-minute period preceding the first fight in each recording, and the post-fight epoch as the 10-minute period following the final fight. If the first fight begins within 10 minutes of the start of the recording, the pre-fight epoch becomes accordingly shorter. Similarly, if the recording ends before 10 minutes have passed since the last fight, the post-fight epoch becomes shorter.

### Pectoral point velocity

We restrict our attention to the pectoral point of each organism 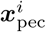 and construct the pectoral point velocity via the centered difference approximation:

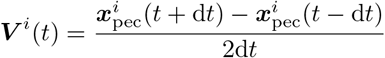

where d*t* = 10^−2^ s is the time step of the recording.

### Projection of velocity onto co-moving coordinates

We introduce a pair coordinate system by defining an axis from the focal individual *i* to another 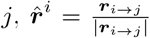, where 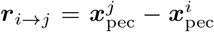, axis of *z* direction 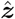, and another orthogonal axis 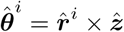. We then project the pectoral point velocity onto these axes:

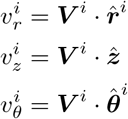

### Cross-correlation function

We compute the time-lagged cross-correlation function from the velocity components of each individual:

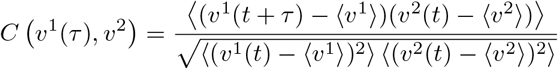

For plots in Fig. 2(C), we compute the cross-correlation function for each epoch and then compute the function’s average, solid line, and standard deviation, shaded area, across recordings.

### Partial information decomposition for two predictor variables and one target variable

We consider the case where we predict a target variable *Y* from two predictor variables *X*_1_ and *X*_2_. Any nonempty subset of predictors can become a *source* of prediction, forming 𝒮 = {{*X*_1_}, {*X*_2_}, {*X*_1_*, X*_2_}. The joint variable is considered another independent source because of the synergistic nature of information (e.g., the XOR operation). Any combination of sources may have non-negative redundant information about the target *Y*, denoted as *I*_∩_(***X***; *Y*), such that *{X*_1_*}* and *{X*_2_*}* have some common information about 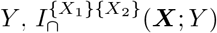, and *{X*_1_*}* and *{X*_1_*}* have self-redundancy, which is equal to the mutual information: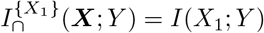. However, if a source *A* is a subset of another source *B*, then 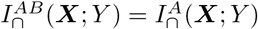, and thus redundant information in any combination of sources from *𝒮* are simplified to four fundamental redundancies 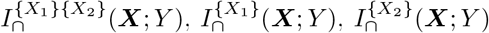, or 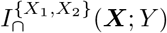. They are ordered such that 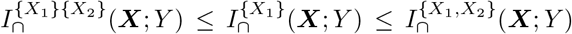 and 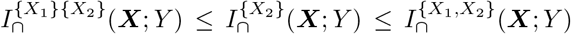.

Therefore,

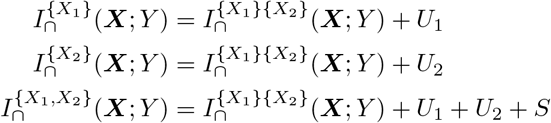

where the unique information in individual variables *U*_1_, *U*_2_, and the synergistic contribution from the joint variable *S* are non-negative. By rewriting 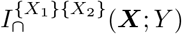 with *R* and self-redundancies with the mutual-information, we obtain the PID for two predictors:

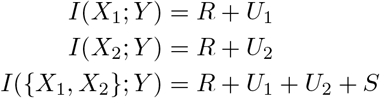

These fundamental redundancies and their ordering can be systematically obtained as *antichains* of sources and the resulting redundancy lattice, respectively, for two or more predictors [8, 18, 19].

The three equations of PID contain four unknowns, and thus one more constrain is required to measure partial informations. In the original PID study from Williams and Beer [8], the redundancy measure is proposed by introducing the *specific information* of an outcome of target *Y* = *y* given the random variable *X*:

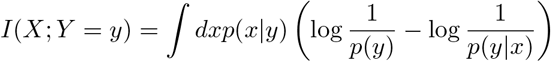

Redundancy is the expected value over all *y* of the minimum specific information that two predictors *X*_1_ and *X*_2_ provide:

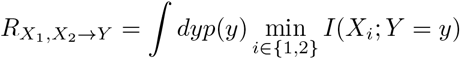

Alternatively, Bertschinger *et al.* [45] defined the unique information, called BROJA unique information, by minimizing the conditional mutual information over a set of hypothetical probability distributions *q* with the constraints that their marginals of each predictor *X_i_* and target *Y* are fixed by the true marginals *p*:

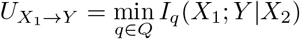

where *Q* = {*q*(*x*_1_*, x*_2_*, y*) *|q*(*x*_1_*, y*) = *p*(*x*_1_*, y*)*, q*(*x*_2_*, y*) = *p*(*x*_2_*, y*)}. PID with BROJA unique information is equivalent to another definition from Griffith *et al.* [46].

### Estimating dynamic information flows from data

To capture changing interactions, we introduce sliding timewindows, and we assume that the underlying joint distribution is Gaussian in each window, Fig. 3(B). The Gaussian predictive information is

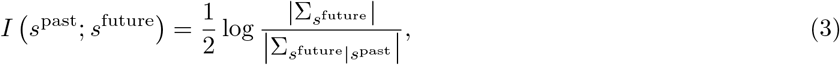

where *|*Σ*|* is the determinant of a covariance matrix Σ, called the generalized variance, measuring the spread of samples [47]. The determinant of the partial covariance 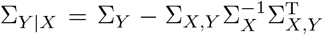 can be simply computed by 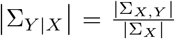 [17]. We estimate the covariance matrix with an adaptive regularization using the OAS estimator [48]. By maximizing the joint information flow 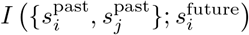, we determine Δ*t*^∗^ = 30 s as the optimal timewindow size for the set of our recordings, Fig. 3(B). For a short time-window, the number of samples is small, and the covariance is highly regularized, resulting small information flow. On the other hand, within long time-windows, distinct linear dynamics appear, correlations decrease, and information flow becomes small.

### Partial information decomposition for Gaussian variables

We use a closed-form solution of redundant information in two Gaussian predictors and a scaler Gaussian target, called “minimum mutual information” [49]:

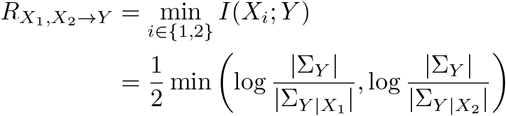

This provides the solution to both PIDs from Williams and Beer [8] and from the BROJA unique information [45] for this Gaussian case. Using the definition of PID provided in Eq. 2 we obtain the other partial informations:

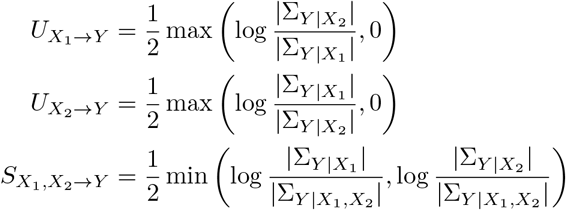

### Relationship between unique-information and correlation fuctions

For PID of predictive information, the opponent-unique information can be expressed as

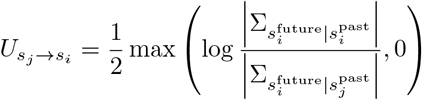

Here, the partial covariance 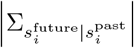 quantifies the conditional variance of 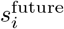 given 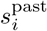, and becomes smaller when 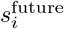 is more predictable from its own past i.e. when autocorrelation is higher. Similarly, 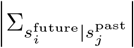 becomes smaller when there is high cross-correlation with 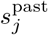. A value of 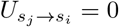 implies

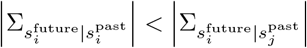

indicating that 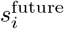 is better predicted from its own past than from the past of *s_j_*, and auto-correlation is higher than cross-correlation. The condition of zero opponent-unique information, observed in our analysis of zebrafish contests, may reflect a generic feature of behavior in which self-predictability exceeds prediction from its environment. We note that this formulation of unique information is valid only under the assumption of a scalar Gaussian target. Alternative formulations, such as incorporating multi-step futures, may reveal non-zero opponent-unique information, as discussed further in the Discussion section.

### Choosing past and future variables

An important step in constructing predictive information is the definition of the past and future variables. Most generally, we can choose the past state as a long history *s*^past^(*t*) = [*x*(*t* −*τ*)*, x*(*t* −*τ* + d*t*)*, …, x*(*t*)] and the future state as a long future trajectory *s*^future^(*t*) = [*x*(*t* + d*t*)*, x*(*t* + 2d*t*)*, …, x*(*t* + *τ*)], where *τ* is the decay time (i.e., *I*(*x*(*t*); *x*(*t* + *τ*)) ∼ 0). In this study, we focus on the one-step future *s*^future^(*t*) = [*x*(*t* + d*t*)] as the closed-form solution of PID for continuous variables only available for a scaler variable [49]. This solution is robust to the dimension of predictor variables, and thus, we adopt *s*^past^(*t*) = [*x*(*t* −*τ* ^∗^)*, x*(*t* −*τ* ^∗^ + d*t*)*, …, x*(*t*)], where the cross-correlation function decays at *τ* ^∗^ (i.e., *C*(*τ*) ∼0 for *|τ| > τ* ^∗^). We use *τ* ^∗^ = 5 s as the length of the past state, thus incorporating the significant cross-correlations, Fig 2(C).

### Covariance estimation

Our target covariance matrix Σ is high-dimensional *p × p*, but the number of samples *n* is limited. In such a case, the sample covariance 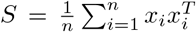 does not minimize the mean squared error *E* {∥*S* −Σ ∥^2^} [48], and the estimation requires a correct regularization. We adopt the oracle approximating shrinkage (OAS) estimator [48]:

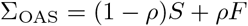

where 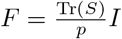 and

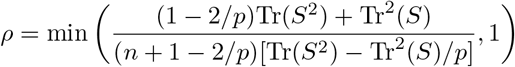

### Computing the average of normalized partial information

We focus on changes in the relative amount of partial informations from each time-window, and thus normalize each partial information by the total information flow, 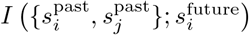. For plots in Fig. 3(C), we first compute the mean of the normalized partial informations within epochs for each recording and obtain the average and SEM across recordings. For plots in Fig. 4, we first align the time by the start and end of the fight epoch and compute the average and SEM of normalized partial informations across recordings at every time point.

---

**FIG. S1:**
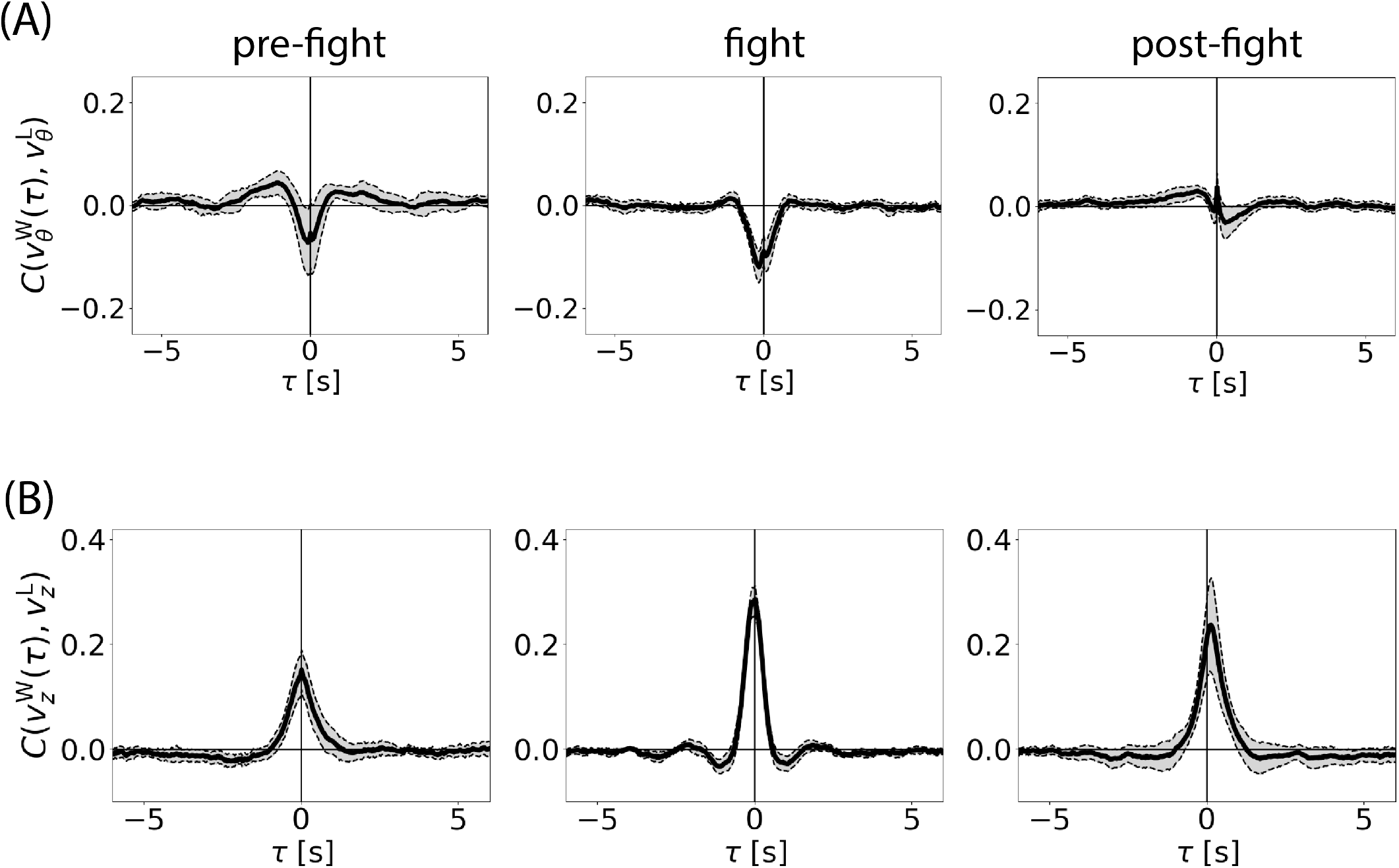
Cross-correlations in *v_θ_* **and** *v_z_* across epochs. (A) We compute the cross-correlation of *v_θ_* between individuals across behavioral epochs, and show the average across 18 experiments. Correlations are higher during the fight phase, reflecting simultaneous directional changes during chases. (B) The average cross-correlation of *v_z_*increases from the pre-fight to fight epoch, indicating that behaviors increasingly involve movement in the *z* direction. The peak of the cross-correlation in the post-fight epoch is slightly shifted toward a positive time lag, suggesting that although the post-fight chase is dominated by winners, movement along the *z*-axis during the chase is led by losers.

**FIG. S2:**
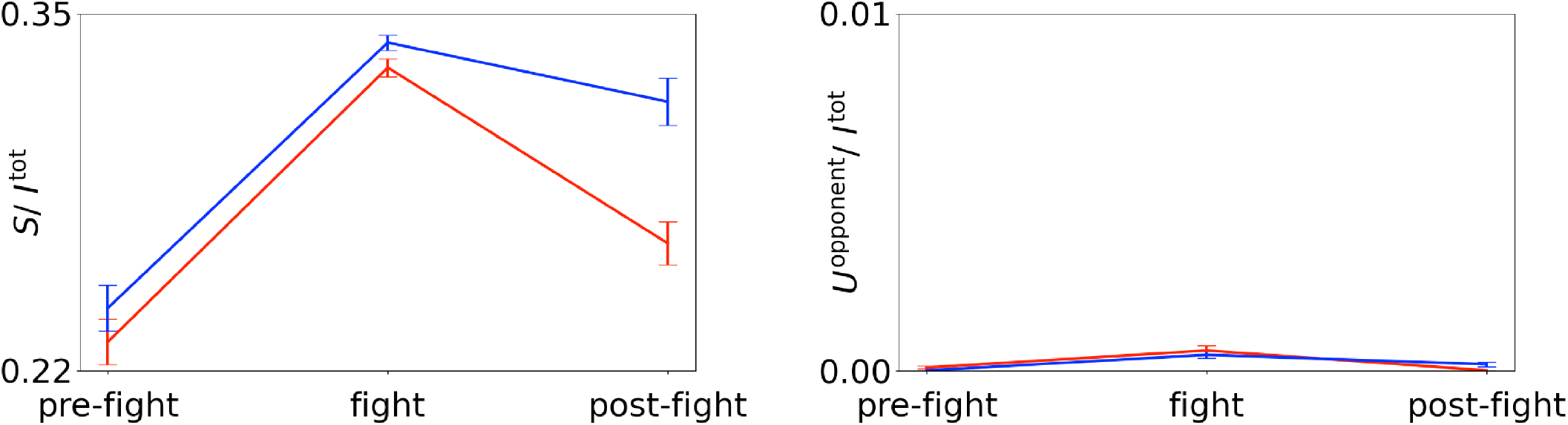
Normalized synergistic information and opponent-self information in *v_r_* across epochs. Synergy follows the same trend of redundancy: high information in the fight epoch and a post-fight asymmetry with higher information in losers than in winners. Opponent-unique information is approximately zero for all epochs as fish movements have inertia, and in their next move in 100 Hz, behavioral elements that uniquely determined by their opponent are negligible.

**FIG. S3:**
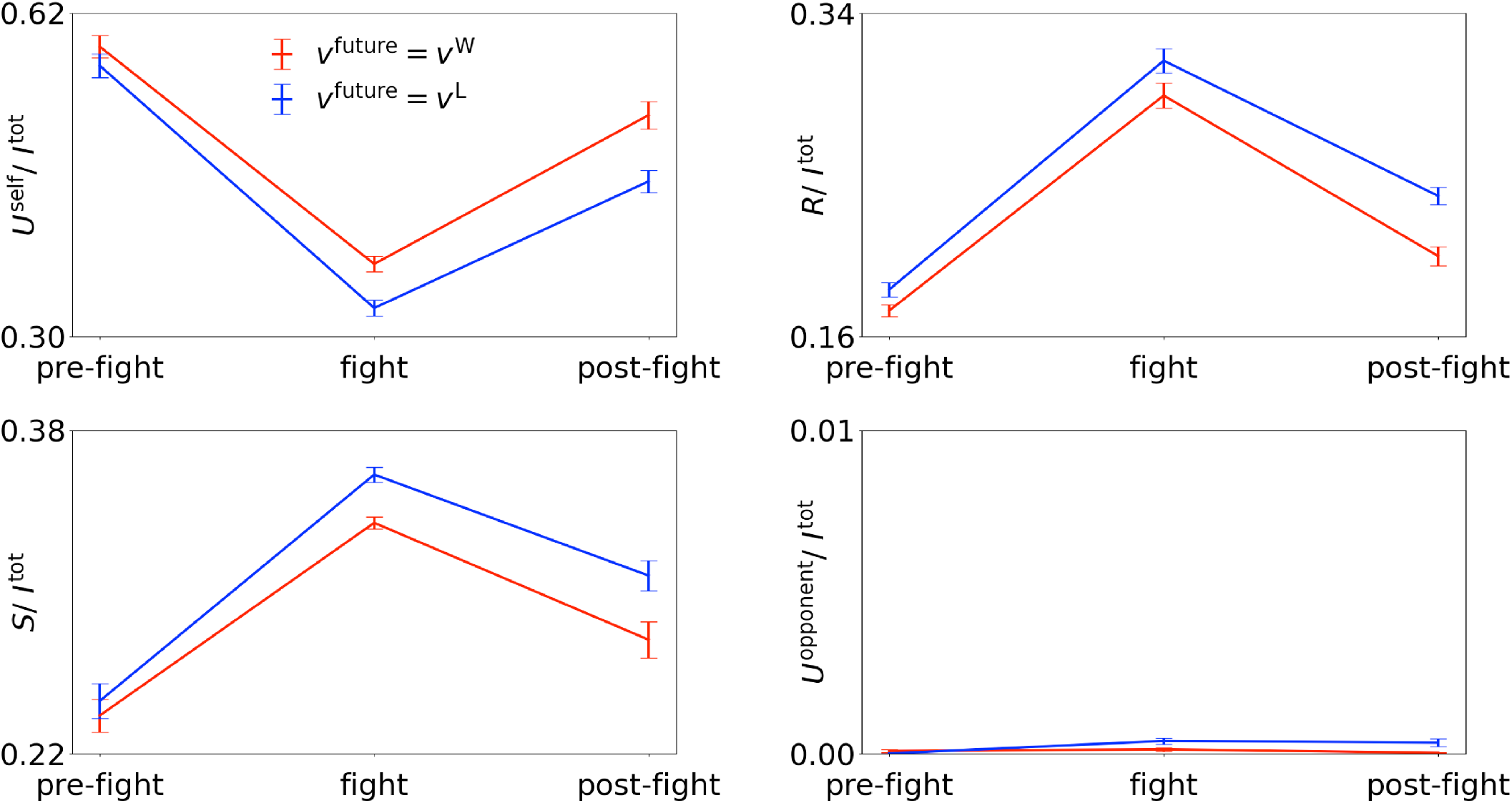
Normalized partial informations in *v_θ_* across epochs. We compute information flows as with *v_r_* and show the average and SEM across experiments.

**FIG. S4:**
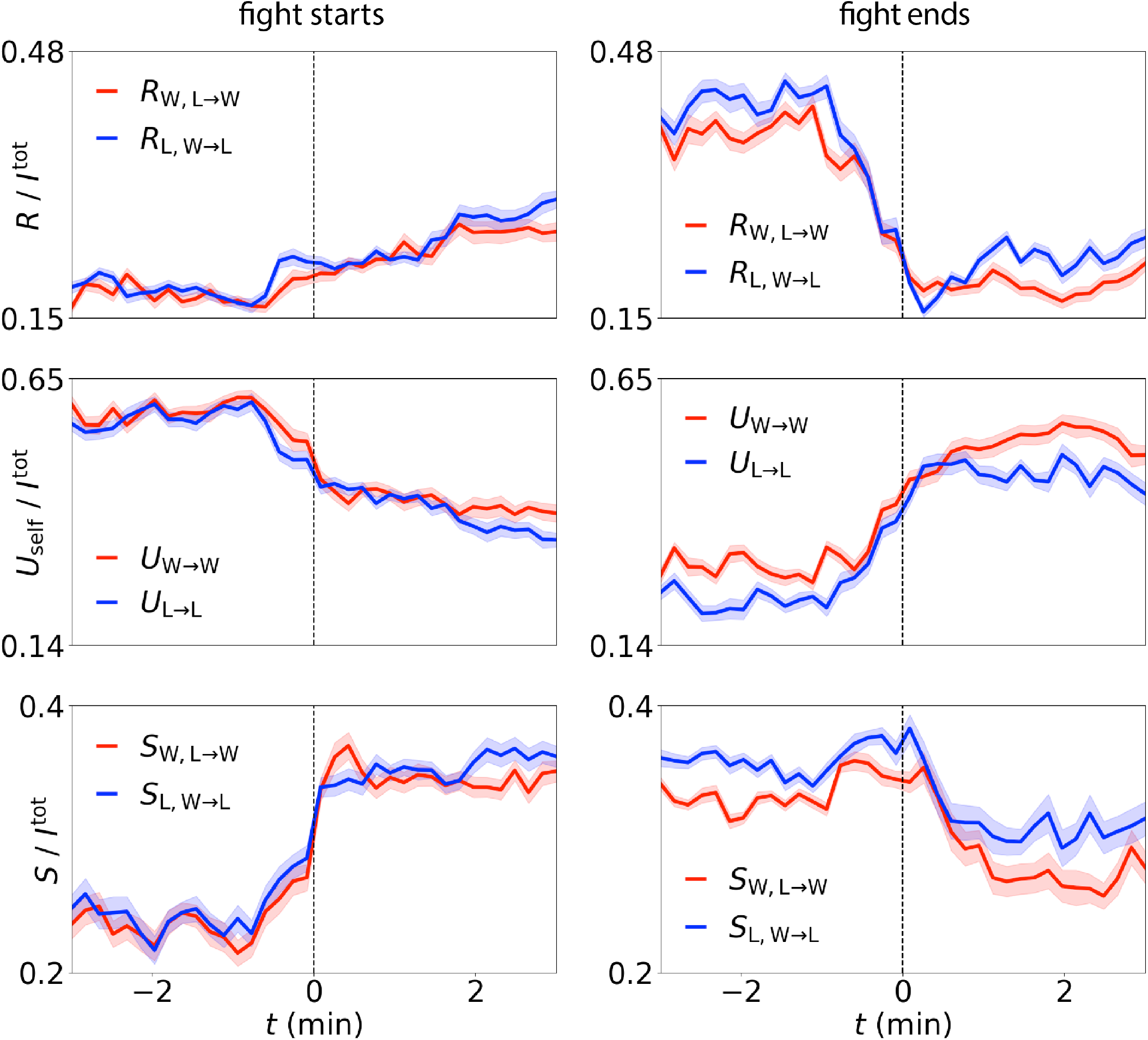
Dynamics of partial informations in *v_θ_* across contest boundaries. (Left) We align by the start of the fight epoch and compute the average and SEM of the fraction of partial information in *v_θ_* across experiments. (Right) We align by the end of fight epoch.

**FIG. S5:**
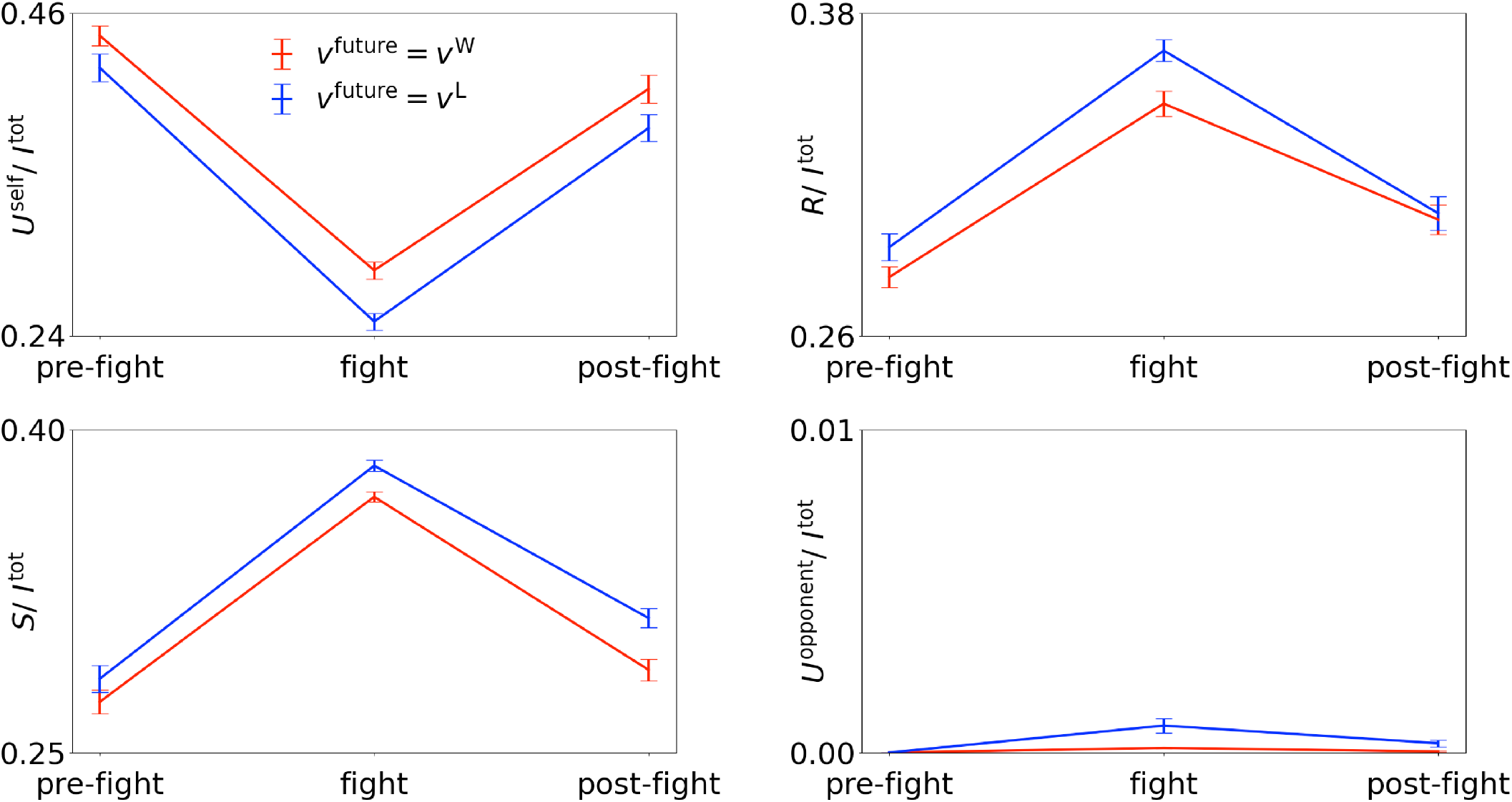
Normalized partial informations in *v_z_* across epochs. We compute information flows as with *v_r_*and show the average and SEM across experiments. In contrast to the results from other coordinates, redundant information in the post-fight epoch is symmetric between winners and losers, implying that although the chase is dominated by winners, movements along the *z*-axis exhibit a similar degree of synchrony.

**FIG. S6:**
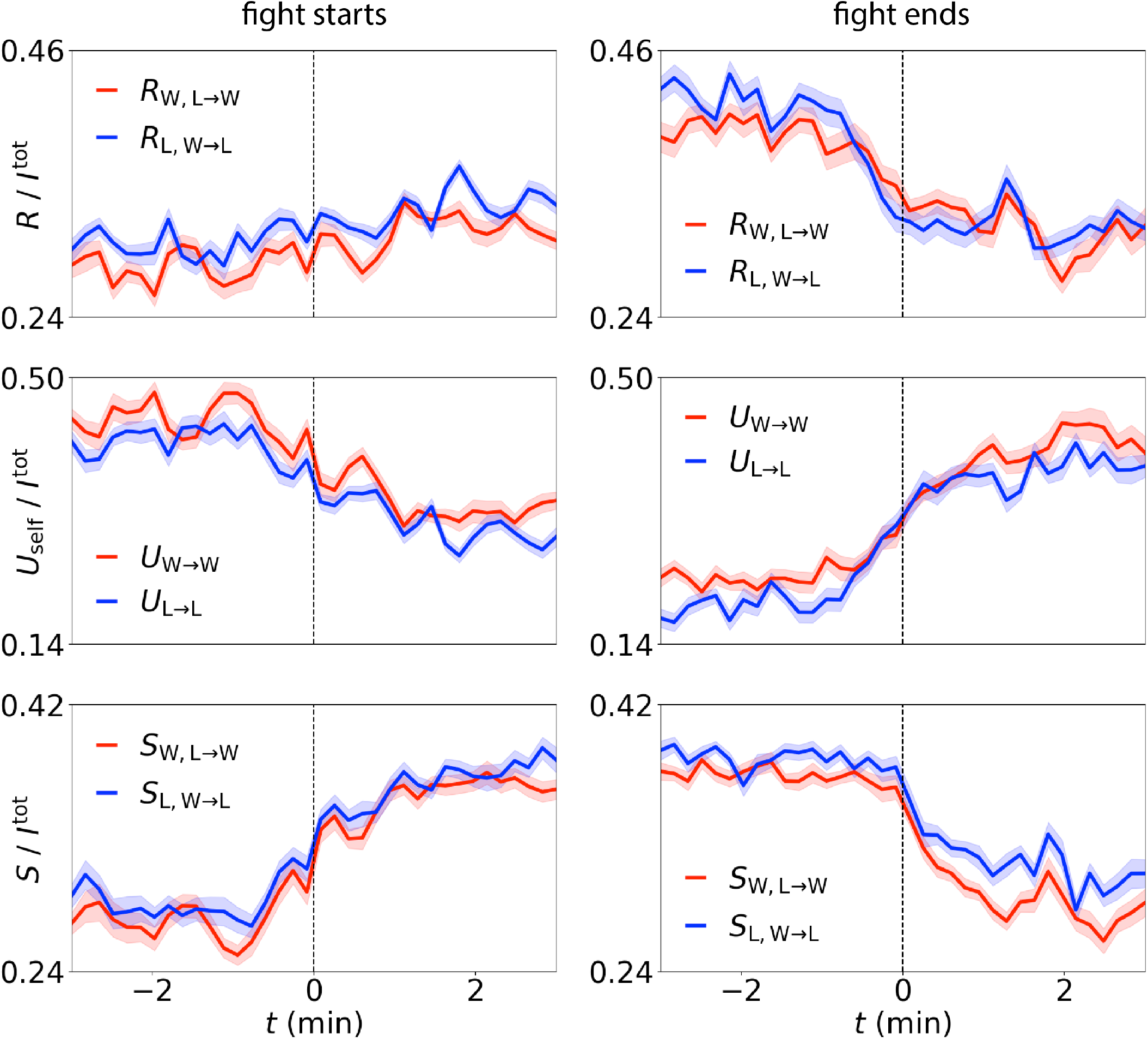
Dynamics of partial informations in *v_z_*across contest boundaries. (Left) We align by the start of the fight epoch and compute the average and SEM of the fraction of partial information in *v_z_*across experiments. (Right) We align by the end of fight epoch.

**FIG. S7:**
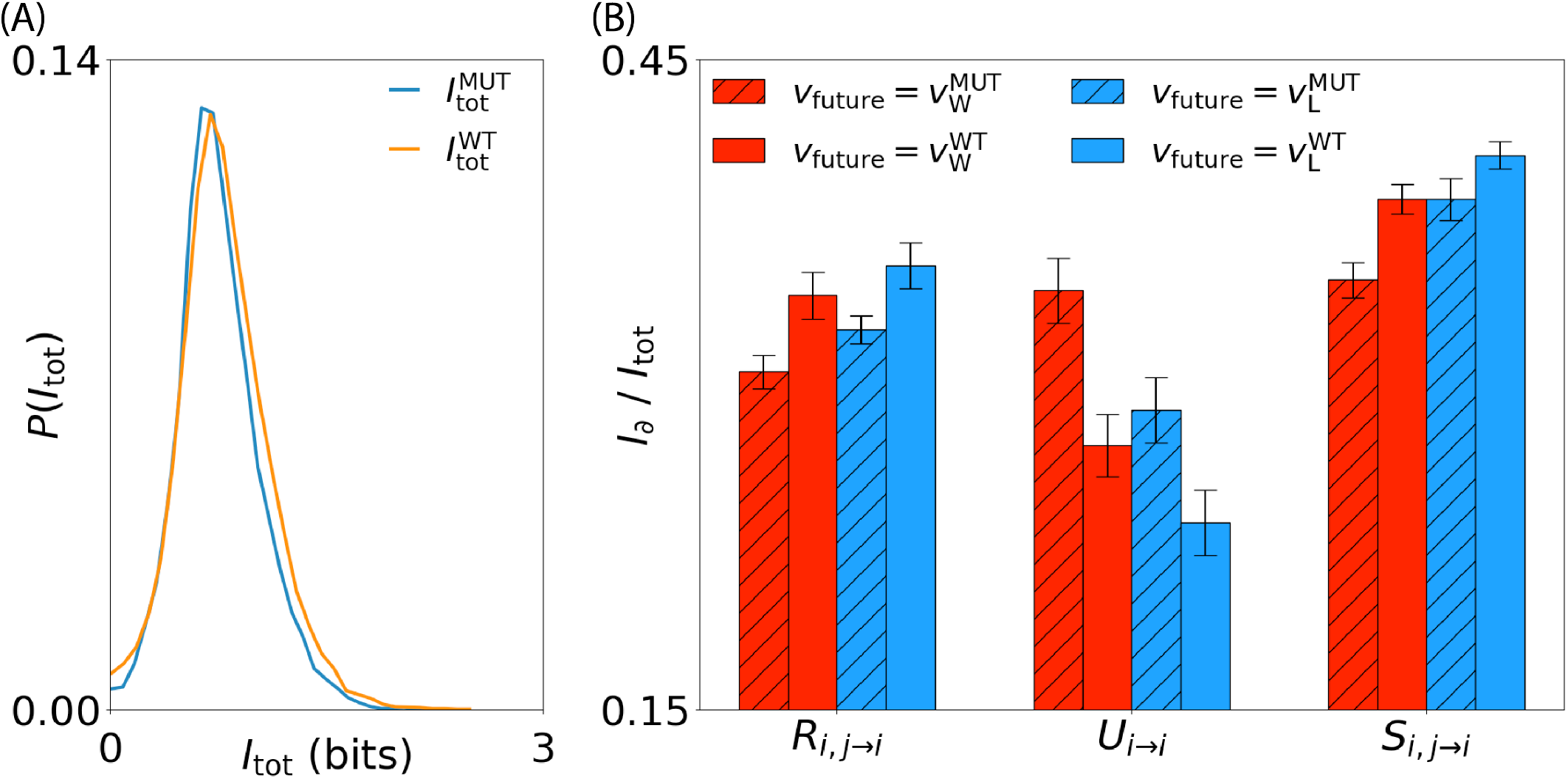
Predictive information in *v_θ_* of *mecp2* mutant pairs and wild-type pairs. We compute predictive information in *v_θ_* from 30 s windows in 5 h recordings of mutant pairs and wild-type pairs. (A) We show the distributions of total information flow across windows (B) We show the normalized partial informations for winners and losers in both mutant and wild-type pairs.

**FIG. S8:**
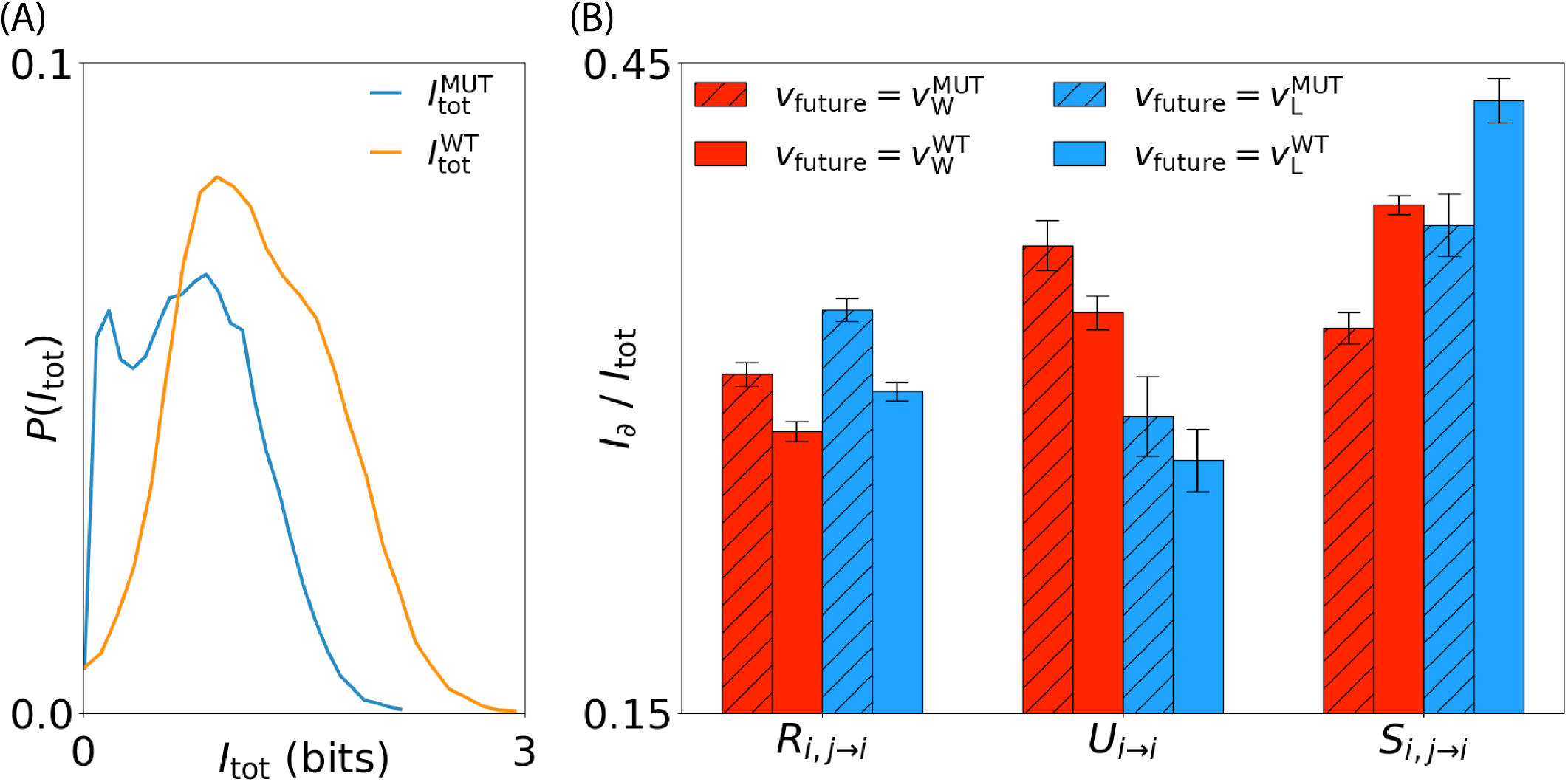
Predictive information in *v_z_*of *mecp2* mutant pairs and wild-type pairs. We compute predictive information in *v_z_*from 30 s windows in 5 h recordings of mutant pairs and wild-type pairs. (A) We show the distribution of total information flow across windows. (B) We show the normalized partial informations for winners and losers in both mutant and wild-type pairs.

SI Movie 1:**Example social behaviors showing characteristic information flows.** We show reconstructed body-points movies with partial information calculated from the corresponding time-window. (Left) Pre-fight interactions with high self-unique information. (Middle) Cyclic attacks with high redundancy and synergy during fight. (Right) Post-fight chase dominated by the winner, showing asymmetric information flows. The movie is directly available here: https://tinyurl.com/r9ase6ep

